# Alpha-synuclein aggregates are phosphatase resistant

**DOI:** 10.1101/2023.11.20.567854

**Authors:** SG Choi, T. Tittle, D. Garcia-Prada, JH Kordower, R Melki, BA Killinger

## Abstract

Alpha-synuclein (αsyn) is an intrinsically disordered protein that aggregates in the brain in several neurodegenerative diseases collectively called synucleinopathies. Phosphorylation of αsyn at serine 129 (PSER129) was considered rare in the healthy human brain but is enriched in pathological αsyn aggregates and is used as a specific marker for disease inclusions. However, recent observations challenge this assumption by demonstrating that PSER129 results from neuronal activity and can be readily detected in the non-diseased mammalian brain. Here, we investigated experimental conditions under which two distinct PSER129 pools, namely endogenous-PSER129 and aggregated-PSER129, could be detected and differentiated in the mammalian brain. Results showed that in the wild-type (WT) mouse brain, perfusion fixation conditions greatly influenced the detection of endogenous-PSER129, with endogenous-PSER129 being nearly undetectable after delayed perfusion fixation (30-minute and 1-hour postmortem interval). Exposure to anesthetics (e.g., Ketamine or xylazine) before perfusion did not significantly influence endogenous-PSER129 detection or levels. In situ, non-specific phosphatase calf alkaline phosphatase (CIAP) selectively dephosphorylated endogenous-PSER129 while αsyn preformed fibril (PFF)-seeded aggregates and genuine disease aggregates (Lewy pathology and Papp–Lantos bodies in Parkinson’s disease and multiple systems atrophy brain, respectively) were resistant to CIAP-mediated dephosphorylation. The phosphatase resistance of aggregates was abolished by sample denaturation, and CIAP-resistant PSER129 was closely associated with proteinase K (PK)-resistant αsyn (i.e., a marker of aggregation). CIAP pretreatment allowed for highly specific detection of seeded αsyn aggregates in a mouse model that accumulates non-aggregated-PSER129. We conclude that αsyn aggregates are impervious to phosphatases, and CIAP pretreatment increases detection specificity for aggregated-PSER129, particularly in well-preserved biological samples (e.g., perfusion fixed or flash-frozen mammalian tissues) where there is a high probability of interference from endogenous-PSER129. Our findings have important implications for the mechanism of PSER129-accumulation in the synucleinopathy brain and provide a simple experimental method to differentiate endogenous-from aggregated PSER129.

**Significance Statement:** Phosphorylated alpha-synuclein (PSER129) was widely regarded as a sensitive, specific marker for pathological aggregates in synucleinopathies until recent data demonstrated that PSER129 is abundant in the healthy mammalian nervous system and results from normal neuronal activity. Differentiating pathological (i.e., aggregated PSER129) and biological (non-aggregated PSER129) has thus become of critical importance to the field. Here, we describe our discovery that aggregated-PSER129 is impervious to enzymatic dephosphorylation. We leverage this discovery to develop a technique (CIAP-PSER129) to detect normal or pathological PSER129 selectively. Our technique allowed us to unambiguously differentiate pathological inclusions in brain regions and mouse models where excessive non-aggregated PSER129 severely limits the sensitivity of aggregate detection. CIAP-PSER129 is nondestructive and compatible with most downstream assays, including mass spectrometry-based peptide identification. These findings have important implications and utility for the synucleinopathy field and may have applicability to other neuropathological proteins (e.g., tau).

## Introduction

Synucleinopathies are age-related neurodegenerative diseases, including Parkinson’s disease (PD), Dementia with Lewy bodies (DLB), and Multiple System Atrophy (MSA), where the prominent neuropathological hallmark is aggregates of misfolded alpha-synuclein (αsyn) protein [1]. Numerous potential mechanisms for αsyn aggregation have been described, but many details remain unclear, particularly within the human diseased brain. Understanding cellular and molecular mechanisms driving αsyn misfolding and aggregation is fundamental for understanding synucleinopathy disease origins and progression.

Fujiwara et al. 2002 identified αsyn phosphorylated at serine 129 (PSER129) within αsyn aggregates from PD, DLB, and MSA brain [2]. Numerous other post-translational modifications (PTMs) associated with αsyn aggregates have been identified [3], with many of these PTMs occurring following αsyn aggregation [4]. In the PD brain, nearly all the αsyn in brain aggregates is PSER129 (∼90%)[2], while endogenous-PSER129 levels in the healthy brain have been estimated to be much less (<5%). However, recent reports suggest that PSER129 occurs during normal neuronal activity [5, 6] and is abundant in specific brain regions [7]. PSER129 may have functional significance by regulating αsyn interactions [8], αsyn-subcellular localization [7], or αsyn-turnover [9, 10]. PSER129 functional significance for disease is unclear, but determining if PSER129 plays a protective[11], neutral[12, 13], or toxic role [14] in the αsyn aggregation processes is important for understanding fundamental synucleinopathy disease mechanisms.

PSER129 detection in the healthy mammalian brain (i.e., endogenous-PER129) has been inconsistent across studies [8, 15–17] with varying abundance, cellular localization, and regional distribution. We observed high variability in the steady-state level of endogenous-PSER129, which could be explained by either sample preparation or biological factors such as neuronal activity [5, 6]. Protein phosphorylation is highly dynamic, cycling between phosphorylation and dephosphorylated forms on short timescales, and as a result, accurate measurement of PTM’s require that biological processes in tissues are rapidly inhibited following death whether by perfusion fixation or other techniques (e.g., flash freezing, small molecule enzyme inhibitors, etc.)[18, 19]. Whether the observed variability in endogenous-PSER129 levels was due to protocol differences or an unknown biological variable is unclear.

Differentiating endogenous-PSER129 and aggregated-PSER129 is crucial for understanding PSER129’s role in αsyn biology and disease. Commercially available PSER129 antibodies have been systematically tested, and EP1536Y (Abcam Cat# ab51253, RRID: AB_869973) has emerged as one of the most specific antibodies for PSER129 [15, 17]. However, this conclusion depends on assay conditions[16], and with abundant endogenous-PSER129, particularly in preclinical models, PSER129 immunoreactivity alone does not alone identify αsyn aggregates. Endogneous-PSER129 labeling in the brain is often incorrectly described as "diffuse," but several brain regions and cell types display strong punctate PSER129 reactivity, which could be mistaken for pathogenic aggregates[16], resulting in false positives. Endogenous-PSER129 may also increase the likelihood of false negatives, as signal amplification methods cannot be fully utilized due to "background" from the endogenous population, which obscures detection[20]. Limited protease digestion enhances specificity for the immunodetection of αsyn aggregates [21], including when using antibodies against PSER129[20]. However, proteases cleave peptide bonds, severely damaging the specimen integrity, limiting practical utility, and impeding downstream assays such as mass spectrometry[22].

Preclinical rodent models have been developed to study αsyn aggregation, including αsyn preformed fibril (PFF) models, which use injections of fragmented synthetically assembled αsyn filaments to "seed" pathology. Mice injected with PFFs into the olfactory bulb (OB-PFF) show robust pathology in select brain regions, including the OB, piriform areas (PA), and entorhinal cortex (EC)[20, 23, 24]. PFF seeding in αsyn transgenic mice (e.g. M83 mice, RRID:IMSR_JAX:004479) can develop more pronounced pathology [25, 26]. The exogenous PFFs in those models remains non-phosphorylated while the seeded endogenous αsyn is phosphorylated[27]. The use of PSER129 as a marker of pathology in preclinical models has been limited by abundant endogenous-PSER129 in certain brain regions (e.g., OB)[16] as well as abnormal hyperphosphorylation before αsyn aggregation[28, 29] that presumably limits detection sensitivity for αsyn aggregates.

## Results

### Rapid perfusion fixation preserves endogenous-PSER129

To investigate why endogenous-PSER129 detection has been variable across studies, we performed delayed perfusion fixation of CO_2_ euthanized mice. To do this, after euthanasia by CO_2_, we immediately removed the animal’s blood by perfusion with PBS and then delayed the time until perfusion with PFA was initiated. We found PSER129 was readily detectable and abundant in brain regions we have previously described [7] in mice rapidly perfused after death (<30s) (Fig. 1A,B). In contrast, 30-min delayed perfusion dramatically reduced PSER129, with only a few PSER129 positive nuclei being detectable in a few brain regions, including the (OB) mitral cell layer (MCL). Similarly, 60-min delayed perfusion resulted in reduced PSER129 staining, with weak reactivity in some cell nuclei of the OB. Quantitative western blots showed that the total amount of αsyn was not significantly altered following delayed perfusion fixation, but PSER129 levels were markedly reduced (F(2,7) = 38.29 , p=0.0002) at 30-min delayed (Optimal vs. 30-min, -97.23% ±0.122), and 60-min (Optimal vs. 60-min, -97.9% ±0.122)(Fig. 1C-E). The distribution of total protein looked similar between all samples (Fig. 1C, left panel).

**Figure 1.**
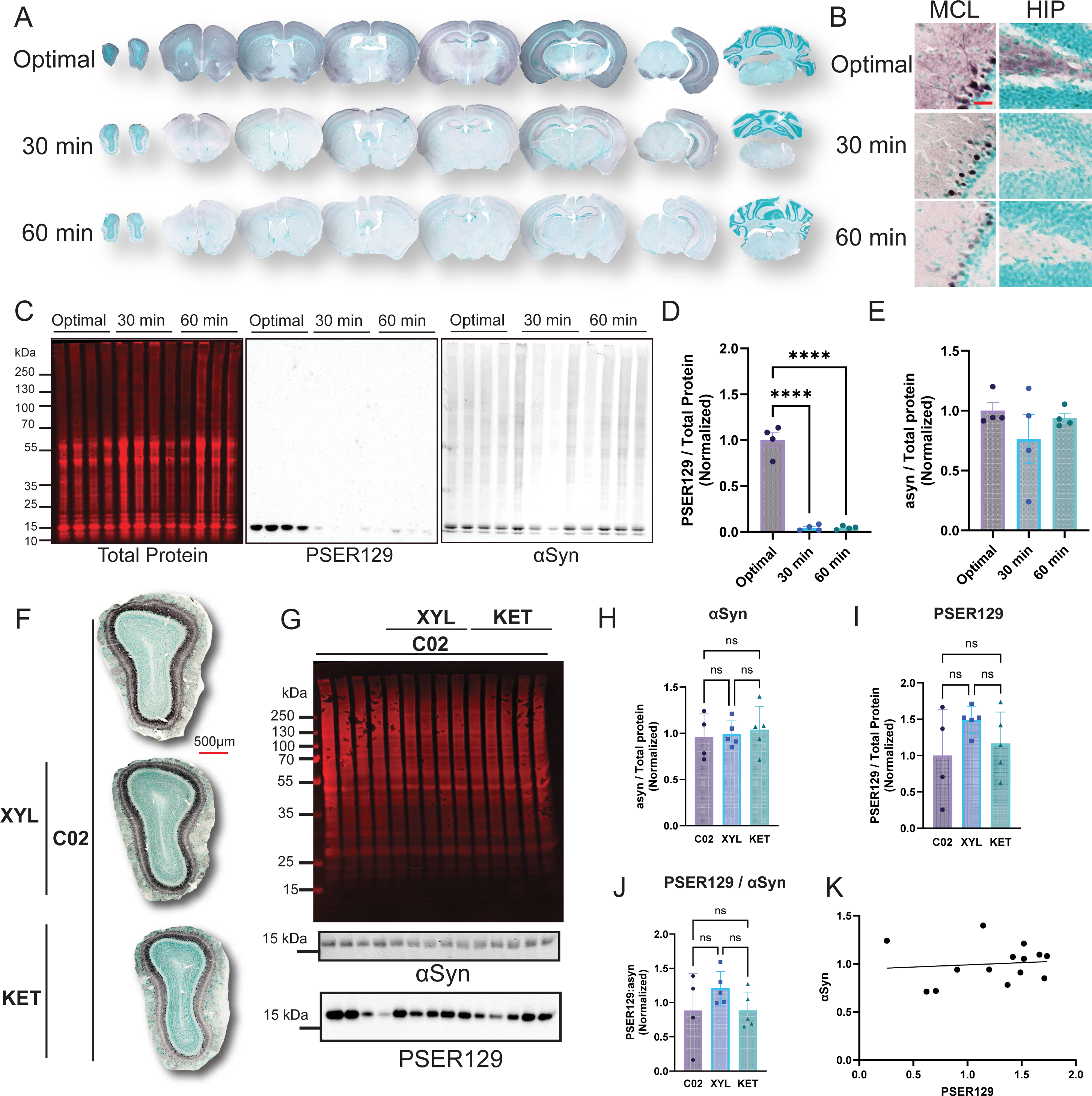
Experimental conditions required for detection of endogenous-PSER129. Rapid perfusion fixation was required for detection of endogenous-PSER129. Mice were euthanized by CO_2_ inhalation, and blood cleared by transcadial perfusion with PBS. Following a delay of 30 or 60 min, mice were then perfused with 4% PFA. (A) PSER129 staining across the neuroaxis. High magnification images of PSER129 staining in the hippocampus (HIP) and the OB mitral cell layer (MCL) using a (B). (C) Proteins extracted from sections across the neuroaxis were resolved by SDS-PAGE and blotted. Blots were probed for total protein (Revert protein stain, Licor), αsyn, and PSER129. Quantification PSER129 (D) and αsyn (E) per protein normalized to the mean of optimal perfusion control. n=4. ****ANOVA, Dunnett’s post-hoc test. p>0.01. Scale bar = 25µm. Anesthetics did not influence PSER129 abundance in the brain. Mice were euthanized by CO_2_ inhalation, and then rapidly fixed by transcadial perfusion with PBS followed by 4% PFA. Ten minutes prior to euthanasia, some animals were exposed to xylazine (XYL, 10mg/kg) or Ketamine (KET, 100mg/kg). (F) PSER129 staining in OB sections. (G) Proteins extracted from sections across neuroaxis were resolved by SDS-PAGE and blotted. Blots were probed for total protein (G, top panel), αsyn, and PSER129. Quantification of total αsyn (H), total PSER129 (I), ratio of PSER129 to αsyn (J), and (K) correlation between αsyn and PSER129 content. No significant differences were found between experimental groups (One-way ANOVA, Tukey’s post-hoc). Total asyn did not correlate with total PSER129 (Pearson Correlation, R^2^ = 0.01045). All sections developed with nickel-DAB Chromogen (black/purple) and counterstained with methylgreen. n=4-5.

### Anesthesia exposure does not influence PSER129 abundance

Ketamine and xylazine are commonly used anesthetics for perfusion and fixation and this procedure could potentially influence endogenous-PSER129 via their effects on neuronal activity[30, 31] or hypothermia [32]. To determine if either anesthetic influenced endogenous-PSER129 detection, we performed perfusion fixation on mice acutely exposed to Ketamine (100mg/kg) xylazine (10mg/kg) 10 min prior to C0_2_-mediated euthanization and perfusion fixation. Results showed that regardless of anesthesia exposure, no significant difference in endogenous-PSER129 content was detected in the mouse OB. IHC detection of PSER129 was similar between groups (Fig. 1F). Quantitative western blot of proteins extracted from PFA-fixed brain sections encompassing the entire brain (Fig. 1G) showed that the amount of αsyn (Fig. 1H), PSER129 (Fig. 1I), or the ratio of PSER129/αsyn (Fig. 1J), did not significantly differ between anesthetic treatments. PSER129 and αsyn quantities were not significantly correlated (Fig. 1K).

### Aggregated-PSER129 selectively resists in situ dephosphorylation in WT mice

To determine if endogenous-PSER129 could be differentiated from aggregated-PSER129, we pretreated brain tissues from OB-PFF-injected mice with calf intestine alkaline phosphatase (CIAP). Without CIAP treatment, PSER129 staining was observed as before [7] throughout the neuroaxis of mice treated with PBS and PFFs (Fig. 2A-C). For OB-PFF mice, in pathology-bearing regions (e.g., OB, EC, AMYG) aggregated- PSER129 was difficult to discern from non-aggregated endogenous-PSER129, except for regions where endogenous-PSER129 levels were relatively low (e.g., GCL). CIAP pretreatment resulted in the abolishment of PSER129 staining in PBS mice, while clear PSER129-positive processes and concentric bodies were observed in selected brain regions of OB-PFF mice, consistent with aggregation seen in the OB-PFF model [20, 23, 33]. CIAP-resistant PSER129 was observed in the MCL of OB-PFF mice, which was obscured entirely by endogenous-PSER129 when CIAP was not used (Fig. 2C). We tested several proteases, but despite having predicted cleavage sites adjacent to αsyn Ser129, they did not wholly abolish PSER129 staining in the PBS mice (Fig. S1). PK can eliminate endogenous-PSER129[20] (Fig. S1), but the treatment is highly destructive, limiting its compatibility with common downstream techniques (LC-MS-based peptide identification). CIAP is nondestructive and effective and should be the preferred approach to differentiate non-aggregated vs. aggregated-PSER129.

**Figure 2.**
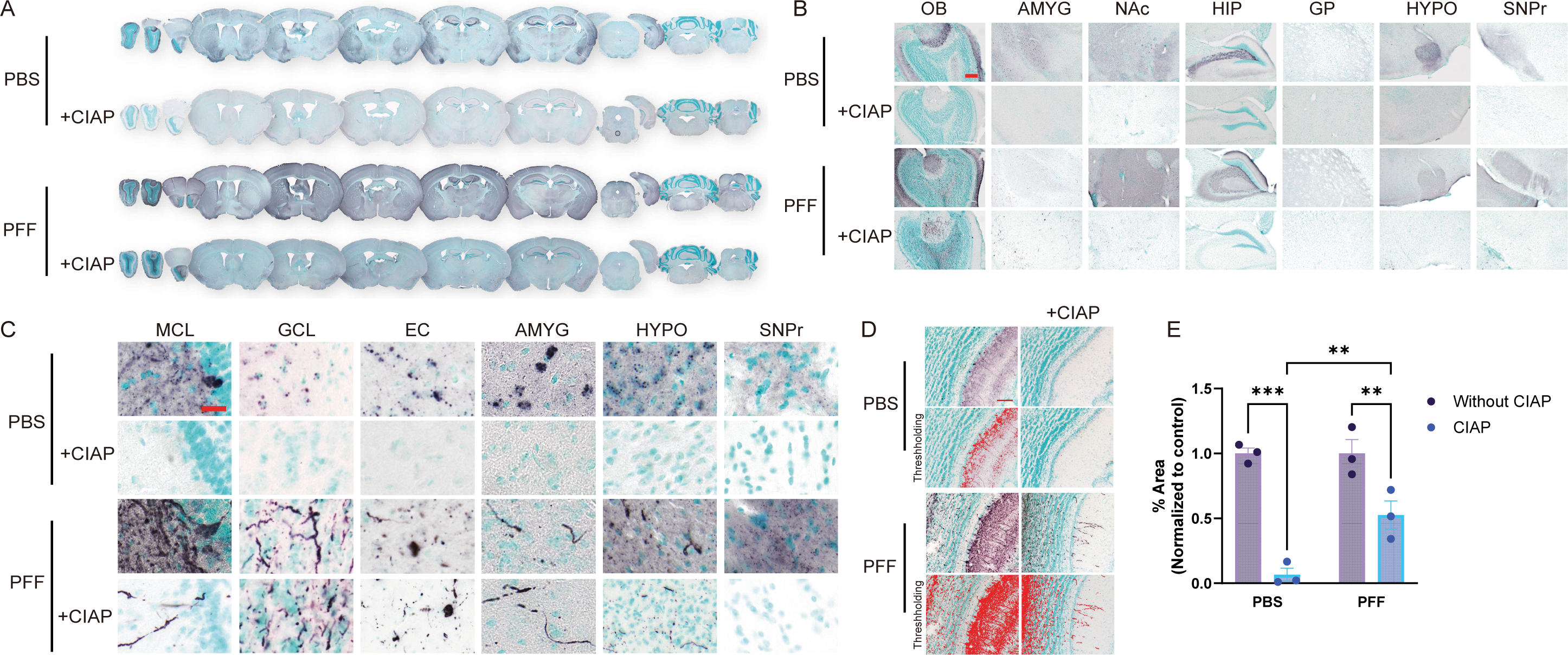
Seeded aggregates in WT mice are CIAP resistant. Mice were bilaterally injected with either PBS or PFFs into the OB GCL. Two months following injection, animals were euthanized with a mixture of ketamine/xylazine followed by transcardial perfusion fixation. Brain sections across the neuroaxis were stained for PSER129 with and without CIAP pretreatment. (A) Whole section images of PSER129 stained brain images across the neuroaxis, with (+CIAP) or without CIAP (-CIAP) pretreatment. (B) Low and (C) high magnification images of select brain regions. (D) Representative images of signal thresholding and (E) subsequent quantification of PSER129 immunoreactivity in the OB GCL. OB = olfactory bulb, AMYG = amygdala, NaAc = nucleus accumbens, HIP = Hippocampus, GP = Globus pallidus, HYPO = hypothalamus, SNPr = substantia nigra pars compacta, MCL = mitral cell layer, GCL = granular cell layer, EC = entorhinal cortex. All sections developed with nickel-DAB Chromogen (black/purple) and counterstained with methylgreen. n=3. Scale BAR for B = 100µm, for C = 25µm, D = 50 µm.

Thresholding was performed on OB images (Fig. 2D), and the PSER129 area was calculated (Fig. 2E). Overall, a significant decrease in PSER129 was observed after CIAP pretreatment (F(1,4) = 131.2 , p=0.0003). Furthermore, following CIAP treatment, PSER129 was higher in PFF-treated mice than in PBS-treated mice (p=0.0045). In contrast, without CIAP, no significant differences were detected between PBS and PFF treated mice.

### Aggregated-PSER129 selectively resists in situ dephosphorylation in M83 mutant mice

Mice overexpressing human A53T αsyn under the prion promoter (i.e., M83) are a common model for studying αsyn aggregation[34] as they accumulate PSER129 throughout the brain after ∼ three months of age[16, 28] but do not develop widespread filamentous αsyn aggregates until around ten months of age with the brain stem being most severely affected [26, 34]. M83 mice injected with extracts from diseased brains or αsyn PFFs develop pathology within a few months of the injection and have an accelerated disease phenotype[26]. Because the M83 mouse brain has high quantities of PSER129 prior to aggregation, differentiation between aggregated-CIAP and endogenous-PSER129 is challenging and can impede accurate assessment of aggregation in this model.

Here we tested whether CIAP treatment could differentiate abundant nonaggregated-PSER129 from aggregate-PSER129 in homozygous M83 mice injected with PFFs unilaterally into the OB. Results showed that without CIAP, ubiquitous PSER129 reactivity was observed in both the injected and noninjected hemispheres (Figure 4A). In the OB and PA, PSER129 reactivity was similar, with a few dysmorphic neurites being observable (Figure 3A, B). However, following CIAP pretreatment, the PSER129 reactivity was restricted to the GCL of the injected OB (Figure 3A). In PA-containing sections just posterior to the OB injection site, diffuse weak PSER129 reactivity was observed, with asymmetrical PSER129 signal in the PA of the injected hemisphere (3A, red outline). High magnification images revealed that following CIAP non-diffuse PSER129 reactivity could be observed only in the GCL and PA of PFF-treated mice (3B). Reactivity was strong in those regions and morphologically resembled neuronal processes and cell bodies, in contrast to diffuse PSER129 in PBS treated mice, which lacked PSER129 reactivity.

**Figure 3.**
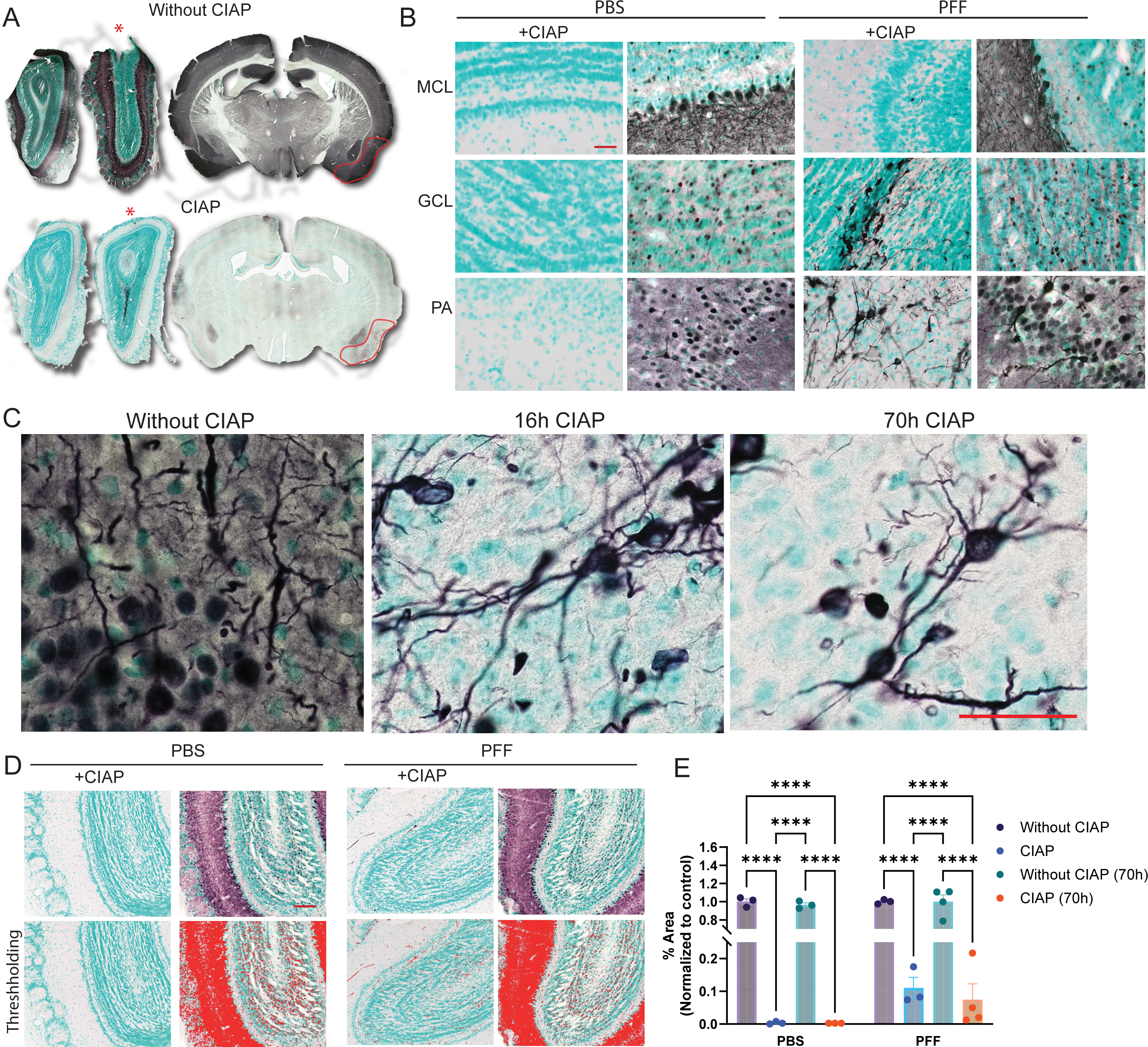
Seeded aggregates in M83 mice are CIAP resistant. M83 mice were unilaterally injected with αsyn PFFs into the OB GCL. 6 months following injections mice were perfusion fixed. (A) PSER129 staining in the brain or M83 mouse injected with PFFs without or without CIAP treatment. Red star denotes the OB injected with PFFs. Red line annotates the PA of the PFF injected hemisphere. (B) High magnification images of CIAP treatment tissue sections from M83 mice injected with either PBS or PFFs. (C) High magnification images of PA from mouse brain sections treated with CIAP for either 16h or 70h. (D) Representative thresholding of images prior to quantification. (E) Quantification of PSER129 signal in the OB. **** Fisher’s LSD, p<0.0001, n=3-4. Scale bar for B,C, and D = 50 µm.

**Figure 4.**
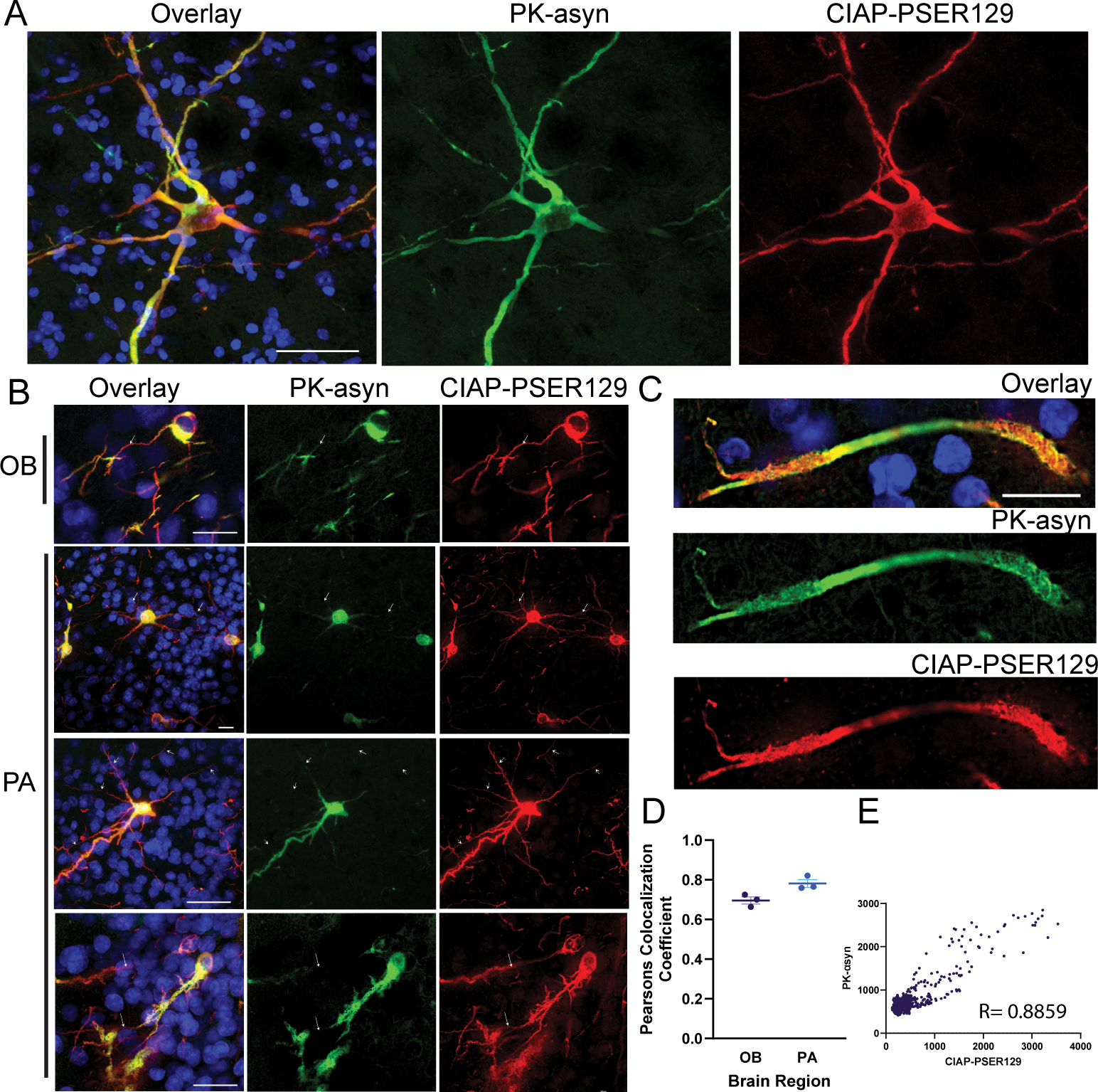
CIAP-resistant PSER129 partially colocalizes with PK-resistant αsyn. Brain sections from M83 mice unilaterally injected with αsyn PFFs into the OB GCL were multiplex labeled for CIAP resistant PSER129 (CIAP-PSER129, red) and PK resistant αsyn (PK-αsyn, green) and nuclei (DAPI, blue). Representative images show labeling in the brainstem (BS) (A), OB and PA (B). Arrows denote the position of cell processes labeled for CIAP-PSER129 but lack overlapping PK-αsyn. (C) Single plane image of a dysmorphic neurite. (D) Pearson R colocalization coefficients for PK-αsyn and CIAP-PSER129 in the OB and PA. (E) Plot of pixel intensity values for confocal image of CIAP-PSER129 and PK-αsyn in PA. Pearson correlation coefficient (R) shown on plot. n=3-4. Scale bars for A=25 µm, B, C = 20 µm.

To determine the degree of CIAP resistance of apparent aggregated-PSER129, we treated tissues from OB-PFF M83 mice with CIAP for 16h and 70h. We found that PSER129 reactivity remained in PA even after several days (70h) of CIAP exposure (3C). These results suggest that αsyn aggregates are impervious to CIAP-mediated dephosphorylation. We did not test time points greater than 70h. Threshold masking and object area analysis were used to quantify CIAP-resistant and CIAP-sensitive PSER129 (3D) under different CIAP conditions. Results show that in the PBS-injected OB, CIAP pretreatment (16h and 70h) resulted in a total loss of PSER129 immuno-reactivity (3E). Following CIAP, PSER129 staining significantly differed between PBS and PFF-treated mice; in contrast, without CIAP, PSER129 was the same between PBS and PFF (Fig. S3).

### CIAP-resistant PSER129 coincides with PK-resistant **α**syn

We observed CIAP-resistant PSER129 in brain regions consistent with spreading aggregates observed in OB-PFF models[23, 24, 33]. Next, we determined whether CIAP-resistant PSER129 was associated with a separate marker of αsyn aggregates, namely PK-resistant αsyn. To do this, we used a multiplex approach to sequentially label CIAP-resistant PSER129 and then PK-resistant αsyn. Our initial experiments used antibody 211 (Santa Cruz Biotechnology Cat# sc-12767, RRID:AB_628318) to label PK-resistant αsyn (Fig. S4), which revealed an overlap between CIAP-PSER129 and PK-αsyn (Fig. S4A). However, we subsequently confirmed antibody 211 specificity for human αsyn (Fig. S4). Thus, to avoid bias detection of human αsyn, we ultimately used antibody EPR20535 (Abcam Cat# ab212184, RRID:AB_2941889), which reacts to both mouse and human αsyn (Fig. S4B,C).

Results using EPR20535 showed that PK-αsyn overlaps well with CIAP-PSER129 in the OB and PA (Fig. 4). Overlap was most apparent in swollen dysmorphic neurites (4C) and cell bodies (Fig. 4A,B). Distal CIAP-PSER129 positive processes were often weakly labeled with PK-αsyn (Fig. 4B, white arrows). Within dysmorphic neurites (4C) and cell bodies (4B), high magnification images revealed PK-αsyn and CIAP-PSER129 only partially overlap. Some CIAP-PSER129 positive neurites were devoid of any PK-αsyn. Interestingly, within the same neuron, CIAP-PSER129 and PK-αsyn sometimes labeled different cellular compartments (Fig. 4B, white arrows). This suggests that CIAP-PSER129 is specific for αsyn aggregates (i.e., overlaps with PK-αsyn), but also detected ancillary αsyn forms that were sensitive to PK treatment. PK-αsyn and CIAP-PSER129 were strongly, but imperfectly colocalized (Fig. 4D, E)

### **α**Syn aggregates in human synucleinopathy brain resist in situ dephosphorylation

PSER129 is a selective, sensitive marker for pathology in the synucleinopathy brain, as endogenous-PSER129 levels are considered low in the non-synucleinopathy brain. Because we found aggregated-PSER129 was impervious to CIAP-mediated dephosphorylation in PFF models, we tested the hypothesis that αsyn aggregates in the human brain were CIAP resistant. We stained PSER129 in PD (Fig. 5A, C) and MSA (Fig. 5B, C) midbrain with and without CIAP pretreatment. Without CIAP, as expected, we observed abundant Lewy pathology (Fig. 5C, left panels) and Papp Lantos bodies (Fig. 5C, right panels) in the PD and MSA brains, respectively. Following CIAP treatment, we observed similar staining in the PD and MSA brains (Fig. 5C, "+CIAP"). Image thresholding (Fig. 5D) and quantification confirmed no statistically significant difference in PSER129 staining with or without CIAP pretreatment (Fig 5E).

**Figure 5.**
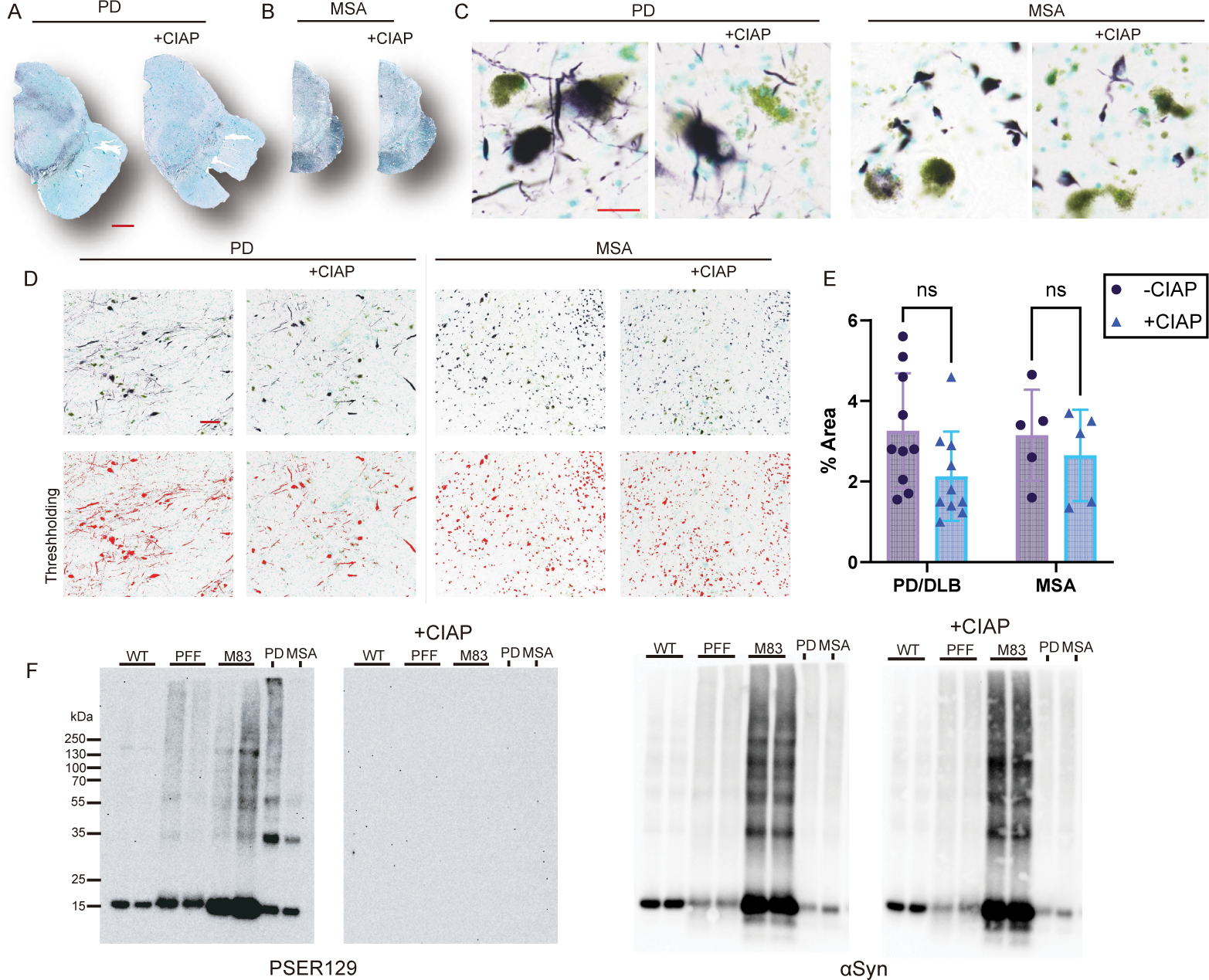
Human brain αsyn pathology is phosphatase resistant. (A) Whole section images of transverse midbrain sections from (A) PD and (B) MSA cases stained for PSER129 with (+CIAP) or without CIAP pretreatment. High magnification images of substantia nigra of (C) PD and MSA. (D) Representative thresholding of images prior to quantification. (E) Quantification of staining in PD and MSA brain with or without CIAP. (F) Proteins extracted from tissues that either did or did not have αsyn aggregates. Proteins were separated by SDS-PAGE and blotted onto PVDF membranes which were incubated with and without CIAP. Blots were probed for both PSER129 and αsyn. All sections developed with nickel-DAB Chromogen (black/purple) and counterstained with methylgreen n=5-10. Scale bars for A and B = 2mm, for C = 25 µm, for D = 150 µm.

### Denaturation abolishes aggregated-PSER129 CIAP resistance

αSyn conformational transition to aggregates imparts resistance to proteases. PFF seeded (Fig. 2-4) and bona fide αsyn aggregates (Fig. and 5A-E) resist CIAP-mediated dephosphorylation in situ. To determine if the observed effect was due to the conformation of aggregates, we denatured the brain samples, separated the proteins by WB, and exposed blotted denatured proteins to CIAP. We did this for proteins from untreated WT mice, OB-PFF WT mice, M83 mice, PD midbrain, and MSA midbrain.

Results show that in the denatured samples, CIAP abolished PSER129 immunoreactivity in non-aggregate containing samples (i.e., untreated WT) and aggregate-containing samples (i.e., OB-PFF, M83, PD, and MSA) (Fig. 5F). CIAP pretreatment abolished the reactivity of monomeric (14kDa) and high molecular weight species (>30kDa). This shows that conformation imparts resistance to CIAP, analogous to protease-resistance.

## Discussion

CIAP pretreatment will help differentiate endogenous- vs. aggregated-PSER129 in preclinical animal models. Protease pretreatment (e.g., PK, Trypsin, GluC) could also be used (Fig. S2), but those treatments are destructive, in particular, PK, and can impede downstream assays (e.g., peptide identification by mass spectrometry, IHC)[21] as well as alter tissue integrity. For this reason, CIAP pretreatment might be a particularly valuable approach to increase specificity for aggregated-PSER129 in multiple assay platforms. Endogenous-PSER129 is CIAP sensitive, and aggregated-PSER129 is CIAP insensitive, this simple observation might help differentiate the two pools in future studies.

CIAP-PSER129 may detect early changes in αsyn conformation. CIAP-resistant PSER129 showed high, but not complete, overlap with PK-resistant αsyn (Fig. 4), demonstrating that CIAP-PSER129 was detecting αsyn aggregates and other structures. Some evidence suggests that PSER129 is an early event in the aggregation process[4, 35]. We speculate that CIAP-PSER129 labeling of the non-dysmorphic neurites of aggregate-containing neurons are early aggregate structures that have not yet become PK resistant. CIAP is a massive protein compared to PK (140kDa and 28.9kDa, respectively), and therefore CIAP enzyme may be sterically hindered even for smaller αsyn aggregates that are sensitive to PK. Indeed, disease-causing αsyn mutations impair PSER129 reversibility prior to overt aggregation [36]. Therefore, CIAP-PSER129 may be capable of detecting minute abnormal αsyn conformations (i.e., misfolding) prior to larger PK resistant aggregate formation. Future studies are needed to examine this possibility.

Our previous work found widespread regionally specific endogenous-PSER129 in the mammalian brain [7]. Here we explain why PSER129 content fluctuated greatly between specimens, with some brain specimens (e.g., rodent and human) showing little or no endogenous-PSER129. We also explain the "all-or-none" effect, where endogenous-PSER129 abundance was proportional across the entire neuroaxis for any rodent brain. Both effects were likely due to differences in the rapidity and completeness of PFA fixation and not because of a global biological process driving endogenous-PSER129 fluctuations in the brain. Rapid (<5 min) complete perfusion fixation was a critical parameter for preserving endogenous-PSER129, similar to what has been observed with other phosphoepitopes[18]. Therefore, studies looking at endogenous PSER129 should ensure proper fixation, and animals where perfusion failed or immersion fixation was used, should be eliminated from analysis. Although not tested here, timely flash freezing or lysis in phosphatase inhibitor buffer can also preserve endogenous-PSER129[5, 17], but investigators should consider any prolonged delay in brain removal, as this may impact endogenous-PSER129 detection.

Our findings suggest an alternative explanation for the long-observed apparent enrichment of PSER129 for αsyn aggregates. Previous studies demonstrated rapid PSER129 dephosphorylation in tissue lysates [2], but here we found endogenous PSER129 (i.e., non-aggregated) is rapidly dephosphorylated intracellularly in the WT mouse brain following death, with PSER129 being undetectable in most brain regions at 1h PMI. In the human brain, endogenous-PSER129 has only been reported in a single OB (i.e., the structure where endogenous PSER129 is most abundant in rodents and primates) from a patient with a short PMI (∼2h). In contrast, endogenous-PSER129 was undetectable in non-synucleinopathy OBs with longer PMIs (>4h). Thus, either the human brain is unique from lower mammals (mouse, rat, non-human primate), and endogenous-PSER129 is not abundant, or endogenous-PSER129 in the human brain has evaded detection due to long PMIs common with human donor brain collection. In support of the latter interpretation, CIAP did not significantly affect PSER129 staining in the human brain (Fig. 6A-E), strongly suggesting that these postmortem tissues do not contain endogenous-PSER129. This is consistent with the scenario that aggregated-PSER129 is preferentially preserved during typical postmortem intervals, and apparent PSER129 enrichment in the synuceinopathy brain may result. We established that genuine human αsyn aggregates (Lewy pathology and Papp-Lantos bodies) resist in vitro dephosphorylation. Therefore, PSER129’s association with aggregates may result from the dephosphorylation of endogenous-PSER129 and preservation of aggregated-PSER129 during the PMI. Several studies have implicated PSER129 as a mediator of αsyn aggregation [11, 14], but our results provide further evidence that, as suggested by others [12, 13, 37], PSER129 accumulation in the synucleinopathy brain may be an epiphenomenon. Future studies should test this hypothesis, as it will help clarify PSER129’s role in synucleinopathies. Furthermore, refining the definition of aggregated-PSER129 (i.e., CIAP resistant) will likely be critical when confirming cases of incidental Lewy pathology based on PSER129 reactivity.

These studies have several limitations. First, we cannot conclude a precise mechanism for the observed loss of PSER129 epitope during the postmortem interval, although enzymatic dephosphorylation or proteolysis likely accounts for our observations. Because we did not observe a reduction in total αsyn with postmortem interval (Fig. 1), endogenous phosphatases are likely responsible, but limited C-terminal truncation or ancillary PTMs near the PSER129 epitope is also possible[17]. Second, these studies do not provide direct evidence of postmortem dephosphorylation of endogenous-PSER129 in the human brain. This point will be critical to test in future studies to understand the relationship between PSER129 and synucleinopathy better.

## Conclusions

In conclusion, aggregated-PSER129 resists enzymatic dephosphorylation. Endogenous non-aggregated-PSER129 can be rapidly dephosphorylated and distinguished from aggregated-PSER129 by pretreatment of specimens with CIAP. CIAP pretreatment increases specificity for detecting αsyn aggregates, and thus, CIAP should be used, particularly in preclinical models. This work helps define and differentiate two distinct PSER129 populations (i.e., non-aggregated and aggregated) in the brain. Furthermore, our results support an alternative hypothesis for the long-observed PSER129 enrichment in aggregates of the postmortem synucleinopathy brain.

## Materials and Methods

### Tissue specimens

C57BL/6J male and female mice (n=24) 4-8 months of age and 4–6-month-old male and female homozygous B6;C3-Tg(Prnp-SNCA*A53T)83Vle/J (M83, RRID:IMSR_JAX:004479)[38] mice (n=8) were used. The Rush University Medical Center institutional animal care and use committee reviewed and approved all animal procedures. Human brain tissues were acquired through the Rush Movement Disorders brain bank and 40-micron floating sections prepared as previously described[39]. See supplementary table 1 for details about human cases.

### PFF injections

WT and M83 mice (n=3) were stereotactically injected with 5 micrograms of human αsyn preformed fibrils (PFFs) into the GCL of the OB (coordinates: coordinates: AP, +5.4 mm; ML, +/− 0.75 mm; DV, −1 mm relative to bregma and dural surface) or PBS and euthanized two months following injection as previously described[20, 33]. WT and M83 mice were injected bilaterally or unilaterally, respectively. All fibril preparations were prepared, sonicated, aliquoted, frozen, and then stored at –80°C. Before injection, aliquots were thawed at 37°C for 3 mins, briefly sonicated, then injected into the OB[24]. Mice were then euthanized for six months following PFF injection.

### Tissue preparation and perfusion

Animal tissues were collected by either anesthetizing with ketamine/xylazine (100 mg ketamine/ kg and 10 mg xylazine/kg), or euthanizing via C0_2_, and then transcrardially perfusing with PBS until the perfusate exiting a small incision in the right atrium was clear. All animals were perfusion fixed, but perfusion fixation was delayed for some animals by clearing blood with PBS and then waiting 30-min (n=4) or 1-hr (n=4) prior to perfusion with 4% PFA. Rapidly perfused mice (n=4, "optimal") were quickly flushed with ice-cold PBS (<30s) and then immediately perfused with ice-cold 4% PFA. All mice in PFF studies were perfusion-fixed by standard methods[40]. Whole brain specimens were removed and post-fixed in 4% PFA at 4°C overnight. M83 mouse brains were immersed for 3 days at 4°C. Brains were then equilibrated in 15% and then 30% sucrose solutions. Brain specimens were frozen and sectioned to 40-micron thickness on a freezing stage microtome. Sections were stored in cryoprotectant solution (30% sucrose, 30% ethylene glycol in PBS) at -20°C.

### Calf Intestine Alkaline Phosphatase (CIAP) treatment

Tissue sections matched for level from each animal were washed three times in dilution media (DM, 50mM Tris-HCl pH 7.2, 150 mM NaCl, 0.05% Triton-X100) and then incubated with 1% Triton X-100 in DM for 10 minutes. Tissues were then incubated with CIAP for 16 hours at 37°C using CIAP buffer (100 mM NaCl, 10 mM MgCl2, and 50 mM Tris-HCl at pH 7.9) containing 30 units of the CIAP enzyme (Promega). Tissues were then washed in DM, and heat-induced antigen retrieval (HIAR) was performed by using sodium citrate buffer (10 mM sodium citrate, 0.05% Tween 20, and pH 6.0) at 85°C for 30 minutes, and then tissues were then cooled to room temperature.

### Immunohistochemistry

Free-floating mouse brain sections were rinsed in dilution media (DM, 50mM Tris-HCl pH 7.4, 150mM NaCl, 0.5% Triton-X100) and incubated in peroxidase quenching solution (0.3% hydrogen peroxide, 0.1% sodium azide) containing blocking buffer (3% goat serum, 2% BSA, 0.4% Triton X-100 in DM) for 1 hour at room temperature. Sections were washed in DM and incubated with PSER129 antibody (Abcam, "EP1536Y", 1:50,000 diluted in blocking buffer) overnight. The next day, tissues were washed and incubated with biotinylated anti-rabbit antibody (Vector Laboratories, dil. 1:200 in blocking buffer) for 1 hour at room temperature, followed by rinsing in DM. The sections were then incubated with prepared elite avidin-biotin complex (ABC) reagent (Vector Laboratories) for 75 min at room temperature. For tyramide signal amplification (TSA) sections were washed in DM and borate buffer (0.05M Sodium Borate, pH 8.5). Samples were then incubated with TSA reaction buffer (1 µg / mL biotinyl tyramide, 0.003% hydrogen peroxide in borate buffer) for 30 minutes and at room temperature. Following the TSA reaction, samples were rinsed in DM and heated to 80°C in citrate buffer for 30 minutes, cooled, and incubated with ABC reagent for 75 minutes at room temperature. Some samples were stained without TSA. Tissues were developed using a standard nickel-enhanced 3,3’-diaminobenzidine (DAB)-imidazole protocol and rinsed with sodium acetate buffer (0.2 M Imidazole, 1.0 M sodium acetate buffer, pH 7.2) and PBS before mounting on glass slides. Sections were counterstained with methyl green, dehydrated, cleaned with xylenes, and cover-slipped with cytosol 60 (Fisher Scientific).

### Multiplex tyramide labeling

Detailed multiplex fluorescent tyramide (FT) labeling protocol is available at protocols.io (dx.doi.org/10.17504/protocols.io.yxmvme7zng3p/v1). We specifically adapted this protocol to label CIAP-resistant PSER129 and PK-resistant αsyn simultaneously. To do this, floating sections were first treated with CIAP, as described above. Following CIAP treatment, endogenous peroxidases were quenched and tissues blocked as described above. Tissues were then incubated with EP1536Y (dil. 1:50,000) overnight at 4°. The next day, the sections were washed in DM and then incubated with biotinylated anti-rabbit-igg antibody (Vector Labs) diluted in a blocking buffer for 1h at room temperature. Sections were washed twice with DM and incubated for 75 minutes with diluted ABC reagent (Vector labs). Sections were washed twice with DM and then once with borate buffer. Sections were then incubated in freshly prepared FT working solution (Borat buffer, 0.003% hydrogen peroxide, 5µM CF568) for 30 min at room temperature, protected from light. Sections were then washed twice, and HIAR performed as described above. Sections were then washed in PBS, placed on superfrost plus glass slides, and dried. Slides were then incubated with PK solution (PBS, PK diluted 1:666) for 30 min at 37°C. The tissues on slides were blocked using Bloxall endogenous blocking solution (Vector Laboratories) for 10 minutes. After rinsing in DM, the slides were incubated with antibody EPR20535 (dil. 1:20,000) overnight at 4°C. The following day, the tissues were washed three times in DM and incubated with a biotinylated secondary anti-rabbit antibody (Vector Labs) for an hour, washed in DM, and incubated with prepared diluted ABC reagent (Vector Labs) for 75 min. The tissues were washed in DM three times and incubated with FT working buffer (Borat buffer, 0.003% hydrogen peroxide, 5µM CF488) for 30 minutes at room temperature. The tissues were washed in PBS, and counterstained with DAPI (Sigma-Aldrich, 1:2000). Slides were covered with #1.5 glass coverslips using FluoroShield mounting medium (Sigma-Aldrich).

### Protein extraction

Proteins were extracted from free-floating PFA fixed tissues sections as previously described [7, 39]. Briefly, sections were rinsed in DM several times, collected into a 1.5mL Eppendorf tube, and then heated to 98°C in reversal buffer (0.5M Tris-HCl pH 8.0, 5% SDS, 150mM NaCl, 2mM EDTA) for 30 min. Samples were vigorously vortexed and heated for an additional 15 min. After cooling to room temperature, samples were centrifuged at 22,000 x g for 20 minutes at room temperature. The supernatant was carefully collected, and a BCA assay was performed on 0.5µl of extract to determine protein concentration.

### Western blotting

15-20 µg of protein were separated using 4-13% Bis-tris gels, blotted onto methanol-activated polyvinylidene difluoride (PVDF) membrane, post-fixed with 4% PFA for 30 min, and then allowed to dry completely. Blots were then reactivated in methanol and stained for total protein (Licor). For CIAP treatment, blots were blocked in 2% (w/v) polyvinylpyrrolidone (PVP) for 1h, and then incubated with CIAP as described above.

Blots were then rinsed with TBST (20 mM Tris-HCl pH 7.6, 150 mM NaCl, 0.1% Tween-20) and placed in blocking buffer (TBST and 5% BSA or 5% dry milk) for 1Lh at room temperature. Blots were incubated overnight with PSER129 diluted 1:50,000 or SYN1 diluted 1:2000 in blocking buffer. Blots were then washed and incubated with anti-rabbit HRP conjugate (dil. 1:20,000, Invitrogen), anti-mouse HRP conjugate (dil. 1:6,000, Cell signaling), or anti-mouse IRDye 680RD (dil. 1:20,000 Li-Cor) diluted in blocking buffer for 1 h. Membranes were washed again in TBST and imaged using enhanced chemiluminescence (ECL) substrates (Bio-Rad, product # 170-5060 or ThermoFisher Scientific, product #38554) with a Chemidoc imager (Bio-Rad) or odyssey M imager for fluorescence (Li-Cor).

### Microscopy and imaging

Prepared slides were imaged with Nikon A1 inverted microscope using a 10X, 20X or 63X objective. Whole section scans were acquired with either the Nikon A1 microscope with 10X objective, or the Odyssey-M whole slide imaging functionality. Whole section scans were imported into adobe photoshop for downsizing, cutting, auto-color balancing, and auto-brightness adjustments. Edited images were imported into adobe illustrator for arrangement and final presentation in figures.

### Quantification of IHC

Brightfield images were captured with an inverted confocal microscope equipped with a 20X objective (Nikon A1R). Annotation of each tissue section was conducted within a bounding box of 2000×2000 pixels for mouse tissues and 2863×2454 pixels for human tissues. Manual RGB-based color thresholding was used for mouse and for human tissues NIS-elements (version 5.10.01, https://www.microscope.healthcare.nikon.com/products/software/nis-elements, RRID:SCR_014329) auto-thresholding algorithm was used. The percentage area of the thresholded signal was exported and normalized to average value of non-CIAP treated tissues.

### Data analysis and figures

DAB-stained tissues were quantified using Nikon Elements software as described above. Western blot quantification was performed using both Licor Emperia studio (version 3.0, https://www.licor.com/bio/empiria-studio/download-empiria) and ImageJ (version 1.54h, https://imagej.net/ij/download.html, RRID:SCR_003070).

Statistical analysis and graphing were performed using GraphPad Prism (Version 10.2.0, https://www.graphpad.com/, RRID:SCR_002798). To compare experimental groups, we used One-way ANOVA with Dunnet’s or Tukey post-hoc tests. Pearson correlation was also performed using GraphPad Prism.

## List of Abbreviations

αsyn: alpha-synuclein
PD: parkinson’s disease
MSA: multiple systems atrophy
CIAP: Calf-intestine alkaline phosphatase
PVP: Polyvinylpyrolidone
PSER129: alpha-synuclein phosphorylated at serine 129
DLB: dementia with lewy bodies
PFF: preformed fibrils
OB-PFF: mice with fibrils injected into the olfactory bulb
OB: olfactory bulb
PA: piriform area
GCL: granular cell layer
EC: entrohinal cortex
AMYG: amygdala
MCL: mitral cell layer
SNpr: substantia nigra pars reticulata
PBS: phosphate buffered saline
TBS: Tris buffered saline
TBST: Tris buffered saline with detergent

## Declarations

### Ethics approval

Human brain tissues were used from the Rush movement disorders brain bank with expressed permission by the Rush institutional review board. Animal studies were conducted in accordance with Rush IACUC approved protocol.

### Consent for publication

Not applicable.

### Availability of data and materials

The datasets used and/or analyzed during the current study are available from the corresponding author upon reasonable request.

### Competing interests

The authors have no competing interests to declare.

### Funding

BAK received support from NIH-NINDS award 1R01NS128467 and Michael J. Fox foundation (MJFF) award MJFF-022480. This work was also supported by NINDS Award R21NS109871 and ASAP-024442 (JHK).

### Author contributions

SGC and BAK planned experiments, performed experiments, and wrote the manuscript. TT and DGP performed experiments. JHK and REM wrote/edited the manuscript.

## Acknowledgements

Human brain samples were generously provided by the Rush Movement Disorders Brain Bank. This work was supported in part by Aligning Science Across Parkinson’s [ASAP-024442] through the Michael J. Fox Foundation (MJFF). REM received support from Association France Parkinson.

## Notes

### Competing Interest Statement

The authors have declared no competing interest.

### Summary of Updates

Updated figures with new analysis. New figures 3 and 4. Revised text.

## References

1. Koga, S., et al., Neuropathology and molecular diagnosis of Synucleinopathies. Molecular Neurodegeneration, 2021. 16(1): p. 83.

2. Fujiwara, H., et al., alpha-Synuclein is phosphorylated in synucleinopathy lesions. Nat Cell Biol, 2002. 4(2): p. 160–4.

3. Manzanza, N.O., L. Sedlackova, and R.N. Kalaria, Alpha-Synuclein Post-translational Modifications: Implications for Pathogenesis of Lewy Body Disorders. Front Aging Neurosci, 2021. 13: p. 690293.

4. Mahul-Mellier, A.L., et al., The process of Lewy body formation, rather than simply alpha-synuclein fibrillization, is one of the major drivers of neurodegeneration. Proc Natl Acad Sci U S A, 2020. 117(9): p. 4971–4982.

5. Ramalingam, N., et al., Dynamic physiological α-synuclein S129 phosphorylation is driven by neuronal activity. npj Parkinson’s Disease, 2023. 9(1): p. 4.

6. Ramalingam, N., et al., Dynamic reversibility of alpha-synuclein serine-129 phosphorylation is impaired in synucleinopathy models. EMBO Rep, 2023: p. e57145.

7. Killinger, B.A., et al., Distribution of phosphorylated alpha-synuclein in non-diseased brain implicates olfactory bulb mitral cells in synucleinopathy pathogenesis. npj Parkinson’s Disease, 2023. 9(1): p. 43.

8. Parra-Rivas, L.A., et al., Serine-129 phosphorylation of α-synuclein is a trigger for physiologic protein-protein interactions and synaptic function. bioRxiv, 2022: p. 2022.12.22.521485.

9. Machiya, Y., et al., Phosphorylated alpha-synuclein at Ser-129 is targeted to the proteasome pathway in a ubiquitin-independent manner. J Biol Chem, 2010. 285(52): p. 40732–44.

10. Chau, K.Y., et al., Relationship between alpha synuclein phosphorylation, proteasomal inhibition and cell death: relevance to Parkinson’s disease pathogenesis. J Neurochem, 2009. 110(3): p. 1005–13.

11. Ghanem, S.S., et al., alpha-Synuclein phosphorylation at serine 129 occurs after initial protein deposition and inhibits seeded fibril formation and toxicity. Proc Natl Acad Sci U S A, 2022. 119(15): p. e2109617119.

12. McFarland, N.R., et al., Alpha-synuclein S129 phosphorylation mutants do not alter nigrostriatal toxicity in a rat model of Parkinson disease. J Neuropathol Exp Neurol, 2009. 68(5): p. 515–24.

13. Azeredo da Silveira, S., et al., Phosphorylation does not prompt, nor prevent, the formation of alpha-synuclein toxic species in a rat model of Parkinson’s disease. Hum Mol Genet, 2009. 18(5): p. 872–87.

14. Karampetsou, M., et al., Phosphorylated exogenous alpha-synuclein fibrils exacerbate pathology and induce neuronal dysfunction in mice. Sci Rep, 2017. 7(1): p. 16533.

15. Delic, V., et al., Sensitivity and specificity of phospho-Ser129 alpha-synuclein monoclonal antibodies. J Comp Neurol, 2018. 526(12): p. 1978–1990.

16. Killinger, B.A., et al., Distribution of phosphorylated alpha-synuclein in non-diseased brain implicates olfactory bulb mitral cells in synucleinopathy pathogenesis. NPJ Parkinsons Dis, 2023. 9(1): p. 43.

17. Lashuel, H.A., et al., Revisiting the specificity and ability of phospho-S129 antibodies to capture alpha-synuclein biochemical and pathological diversity. NPJ Parkinsons Dis, 2022. 8(1): p. 136.

18. Wang, Y., et al., Rapid alteration of protein phosphorylation during postmortem: implication in the study of protein phosphorylation. Scientific Reports, 2015. 5(1): p. 15709.

19. Gartner, U., et al., Postmortem changes in the phosphorylation state of tau-protein in the rat brain. Neurobiol Aging, 1998. 19(6): p. 535–43.

20. Walton, S., et al., Neither alpha-synuclein-preformed fibrils derived from patients with <EM>GBA1</EM> mutations nor the host murine genotype significantly influence seeding efficacy in the mouse olfactory bulb. bioRxiv, 2023: p. 2023.08.24.554646.

21. Neumann, M., et al., Misfolded proteinase K-resistant hyperphosphorylated alpha-synuclein in aged transgenic mice with locomotor deterioration and in human alpha-synucleinopathies. J Clin Invest, 2002. 110(10): p. 1429–39.

22. Dau, T., G. Bartolomucci, and J. Rappsilber, Proteomics Using Protease Alternatives to Trypsin Benefits from Sequential Digestion with Trypsin. Analytical Chemistry, 2020. 92(14): p. 9523–9527.

23. Rey, N.L., et al., Transfer of human alpha-synuclein from the olfactory bulb to interconnected brain regions in mice. Acta Neuropathol, 2013. 126(4): p. 555–73.

24. Rey, N.L., et al., alpha-Synuclein conformational strains spread, seed and target neuronal cells differentially after injection into the olfactory bulb. Acta Neuropathol Commun, 2019. 7(1): p. 221.

25. Luk, K.C., et al., Pathological alpha-synuclein transmission initiates Parkinson-like neurodegeneration in nontransgenic mice. Science, 2012. 338(6109): p. 949–53.

26. Luk, K.C., et al., Intracerebral inoculation of pathological alpha-synuclein initiates a rapidly progressive neurodegenerative alpha-synucleinopathy in mice. J Exp Med, 2012. 209(5): p. 975–86.

27. Gribaudo, S., et al., Propagation of alpha-Synuclein Strains within Human Reconstructed Neuronal Network. Stem Cell Reports, 2019. 12(2): p. 230–244.

28. Bencsik, A., et al., Early and Persistent Expression of Phosphorylated α-Synuclein in the Enteric Nervous System of A53T Mutant Human α-Synuclein Transgenic Mice. Journal of Neuropathology & Experimental Neurology, 2014. 73(12): p. 1144–1151.

29. Schell, H., et al., Nuclear and neuritic distribution of serine-129 phosphorylated alpha-synuclein in transgenic mice. Neuroscience, 2009. 160(4): p. 796–804.

30. Mittal, S., et al., beta2-Adrenoreceptor is a regulator of the alpha-synuclein gene driving risk of Parkinson’s disease. Science, 2017. 357(6354): p. 891–898.

31. Cichon, J., et al., Ketamine triggers a switch in excitatory neuronal activity across neocortex. Nat Neurosci, 2023. 26(1): p. 39–52.

32. Planel, E., et al., Anesthesia leads to tau hyperphosphorylation through inhibition of phosphatase activity by hypothermia. J Neurosci, 2007. 27(12): p. 3090–7.

33. Rey, N.L., et al., Widespread transneuronal propagation of alpha-synucleinopathy triggered in olfactory bulb mimics prodromal Parkinson’s disease. J Exp Med, 2016. 213(9): p. 1759–78.

34. Giasson, B.I., et al., Neuronal α-Synucleinopathy with Severe Movement Disorder in Mice Expressing A53T Human α-Synuclein. Neuron, 2002. 34(4): p. 521–533.

35. Sonustun, B., et al., Pathological Relevance of Post-Translationally Modified Alpha-Synuclein (pSer87, pSer129, nTyr39) in Idiopathic Parkinson’s Disease and Multiple System Atrophy. Cells, 2022. 11(5).

36. Ramalingam, N., et al., Dynamic reversibility of alpha-synuclein serine-129 phosphorylation is impaired in synucleinopathy models. EMBO Rep, 2023. 24(12): p. e57145.

37. Buck, K., et al., Ser129 phosphorylation of endogenous alpha-synuclein induced by overexpression of polo-like kinases 2 and 3 in nigral dopamine neurons is not detrimental to their survival and function. Neurobiol Dis, 2015. 78: p. 100–14.

38. Giasson, B.I., et al., Neuronal alpha-synucleinopathy with severe movement disorder in mice expressing A53T human alpha-synuclein. Neuron, 2002. 34(4): p. 521–33.

39. Killinger, B.A., et al., In situ proximity labeling identifies Lewy pathology molecular interactions in the human brain. Proc Natl Acad Sci U S A, 2022. 119(5).

40. Walton, S., et al., Neither alpha-synuclein-preformed fibrils derived from patients with GBA1 mutations nor the host murine genotype significantly influence seeding efficacy in the mouse olfactory bulb. bioRxiv, 2023.

