## Supplemental appendix for "Alpha-synuclein aggregates are phosphatase resistant"

**Supplementary Methods**

*Immunohistochemistry (IHC)*

Free-floating sections were washed in DM and incubated in peroxidase quenching solution containing blocking buffer for 1 hour at room temperature. Sections were then incubated with PSER129 antibody (Abcam, “EP1536Y”, 1:50,000 diluted in blocking buffer) overnight. The next day, tissues were washed and incubated with biotinylated anti-rabbit antibody (Vector Laboratories, 1:200 in blocking buffer) for 1 hour at room temperature, followed by rinsing in DM. The tissues were incubated with prepared elite avidin-biotin complex (ABC) reagent (Vector Laboratories) for 75 min at room temperature and washed in DM and sodium acetate buffer (0.2 M Imidazole, 1.0 M sodium acetate buffer, pH 7.2). Sections were then developed using a standard nickel-enhanced 3,3'-diaminobenzidine (DAB)-imidazole protocol and rinsed with sodium acetate buffer and PBS prior to mounting on glass slides. Tissues were counterstained with methyl green, dehydrated, cleaned with xylenes, and cover-slipped with cytoseal 60 (Fisher Scientific).

*Endoproteinase GluC Treatment*

Tissues matched in position from each animal were rinsed three times in DM. Sections were subjected to protein digestion by incubating with 100 µl of 2X GluC reaction buffer (BioLabs) and 100 µl of stock GluC (BioLabs, 100 ng/µl dissolved in high-purity water) at 37° C for 16 hours.

*Trypsin Treatment*

Tissues underwent protein digestion by incubating with 4 µl of trypsin stock (Sigma-Aldrich, 1 mg/ml dissolved in 1 mM HCl, pH3) diluted in 196 µl of TBS (50 mM Tris-HCl pH 7.6, 150 mM NaCl) at 37° C for 16 hours.

*Proteinase K Treatment*

Protein digestion was performed by incubating the tissues with 1µL of Proteinase K (Thermo Fisher Scientific Inc., ~20 mg/ml) diluted in 199 µL PBS at 37° C for 30 seconds.

*Cleavage site prediction*

Full length mouse alpha-synuclein (>sp|O55042|SYUA_MOUSE Alpha-synuclein OS=Mus musculus OX=10090 GN=Snca PE=1 SV=2 MDVFMKGLSKAKEGVVAAAEKTKQGVAEAAGKTKEGVLYVGSKTKEGVVHGVTTVAEKTKEQVTNVGGAVVTGVTAVAQKTVEGAGNIAAATGFVKKDQMGKGEEGYPQEGILDMPVDPGSEAYEMPSEEGYQDYEPEA) was analyzed using peptide cutter prediction tool software (<https://web.expasy.org/peptide_cutter/>).

*Antigen retrieval*

Following pretreatment with CIAP, floating sections were briefly washed in buffer, heated in antigen retrieval buffer 80°C for 30 minutes, and then allowed to cool to room temperature in antigen retrieval buffer. Several antigen retrieval buffers were tested including citrate buffer without SDS (10 mM Sodium citrate, 0.05% Tween 20, pH 6.0) (“HIAR”) and with SDS (10 mM Sodium citrate, 0.05% Tween 20, pH 6.0) (“0.3% SDS HIAR.” Proteinase K digestion antigen retrieval was also conducted by first mounting and drying sections on glass slides (Superfrost plus slides, Cardinal Health) and the incubating sections with PK (Thermofisher Scientific, Cat#25530049) diluted 1:200 in PBS at 37°C for 30 seconds. Following antigen retrieval IHC was conducted to detect PSER129.

*Multiplex tyramide labeling*

Floating tissue sections from untreated WT and M83 mice were multiplex labeled as described in the main document. Antibody 211 conjugated to HRP (Santa Cruz Biotechnology Cat# sc-12767, RRID:AB_628318) was first labeled with CF569-TF followed by Pan αasyn antibody EPR20535 CF488-TF.

**Supplementary Information**

**Supplementary Figures**

**Supplementary Figure 1**. PSER129 staining of brain sections from OB-PFF mice pretreated with different enzymes. (A) Predicted cleavage sites for digestion of full-length mouse αsyn with glutamyl endopeptidase and trypsin. (B) Whole section images following PSER129 staining using IHC with pre-treatment of CIAP, endoproteinase GluC, and Trypsin for 16 hours. The red boxed regions were enlarged in higher magnification. (C) Enlarged representative images of PSER129 in amygdala and dentate gyrus in PFF-treated mice with pre-treatment of CIAP, endoproteinase GluC, and Trypsin. Scale bars are 50 and 10µm.

**Supplementary Figure 2**. PSER129 immunoreactivity in OB-PFF mice following CIAP pretreatment with different antigen retrieval conditions. (A) Whole sections images PFF-treated mice brain. Tissues were first exposed to CIAP followed by different antigen retrieval conditions, including usual HIAR, 0.3% SDS-containing HIAR, and proteinase K digestion. (B) High magnification images of PSER129 in the boxed regions. Scale bars are 50 and 10µm.

**Supplementary Figure 3.** Additional graphs for Figure 4. Quantification of PSER129 staining in OB sections from M83 mice following different CIAP pretreatment times (A) and without CIAP pretreatment (B). *Two-way ANOVA, F (1, 9) = 6.787, p<0.05).

**Supplementary Figure 4.** CIAP-PSER129 and PK-αsyn. (A) Granule cell layer (GCL) and Ipsilateral piriform area (PA) of OB-PFF-M83 mice multiplex labeled for both CIAP-PSER129 and PK-αsyn using antibodies EP1536Y and 211, respectively. White arrow denotes positions where CIAP-PSER129 and PK-αsyn directly overlap. (B) Brain sections from non-injected M83 mice or WT mice multiplex labeled with two αsyn antibodies, EPR20535 and 211. Images show 211 is only reactive in M83 mice that express human αsyn. In contrast, EPR20535 labeled both WT and M83 mice. This result confirmed 211 is specific for human αsyn and shouldn’t be used if both mouse and human αsyn are to be detected. All images acquired with Nikon A1 confocal microscope.

**Supplementary Table 1.** *Characteristics of MSA and PD/DLB patients.*

| **Sample #** | **^*^Disease** | **Sex** | **Age at death (yrs)** | **^+^Disease Duration (yrs)** | **Hoehn and Yahr Scale** | **Method of fixation** |
| --- | --- | --- | --- | --- | --- | --- |
| 1 | Clinical diagnosis parkinsonism.  Pathological diagnosis MSA. | F | 75 | 2 | 4 | Immersion fixation 7-10 days |
| 2 | Atypical Parkinsonian disorder, probable MSA-P. | F | 65 | 10 | 5 | Immersion fixation 7-10 days |
| 3 | MSA-P | M | 55 | 6 | 4 | Immersion fixation 7-10 days |
| 4 | MSA | M | 67 | 8 | 5 | Immersion fixation 7-10 days |
| 5 | PD primary diagnosis and MSA-P. | M | 60 | 2 | 3 | Immersion fixation 7-10 days |
| 6 | Idiopathic PD. On levodopa, had dyskinesia. | F | 81 | 10 | 5 | Immersion fixation 7-10 days |
| 7 | Lewy body dementia. Cognitive symptoms have dominated over motor/PD symptoms. | M | 73 | 11 | 2 | Immersion fixation 7-10 days |
| 8 | PD/Dementia/Progressive supranuclear palsy. | M | 70 | 5 | 4 | Immersion fixation 7-10 days |
| 9 | PD. | M | 80 | 8 | 4 | Immersion fixation 7-10 days |
| 10 | Dementia. PD secondary. | F | 87 | 3 | 4 | Immersion fixation 7-10 days |
| 11 | PD | F | 87 | 2 | 3 | Immersion fixation 7-10 days |
| 12 | PD.   Subacute dyskinesia due to drug. | F | 89 | 4 | 4 | Immersion fixation 7-10 days |
| 13 | DLB | M | 80 | 1 | 4 | Immersion fixation 7-10 days |
| 14 | PD | M | 84 | 5 | 4 | Immersion fixation 7-10 days |
| 15 | PD.  Cerebral hemorrhage. | M | 80 | 3 | 4 | Immersion fixation 7-10 days |

^*^Except sample #1, all other cases are based on clinical diagnosis. ^+^ The duration of years represents the period from when symptoms were observed/noticed until death, as documented in the clinical records of the Rush Movement Disorder program.
