## Supplemental Figures for "Alpha-synuclein aggregates are phosphatase resistant"

A

| Enzyme | Number of Cleavage sites | Positions of Cleavage sites |
|---|---|---|
| Trypsin | 15 | 6 10 12 21 23 32 34 43 45 58 60 80 96 97 102 |
| Glutamyl endopeptidase | 18 | 13 20 28 35 46 57 61 83 104 105 110 114 123 126 130 131 137 139 |

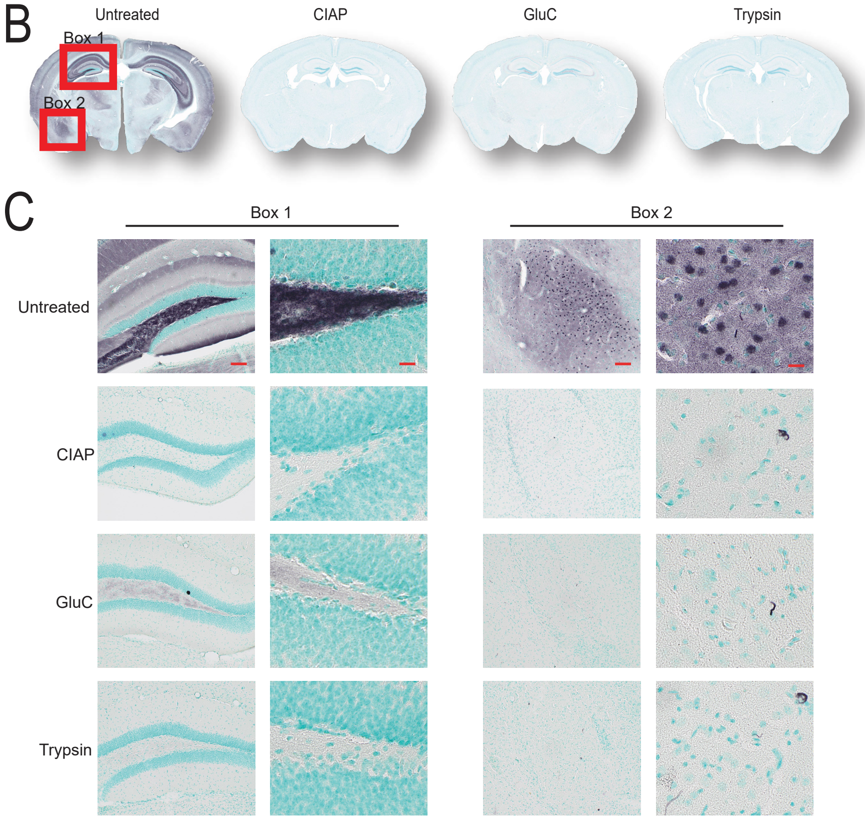


Supplementary Figure 1

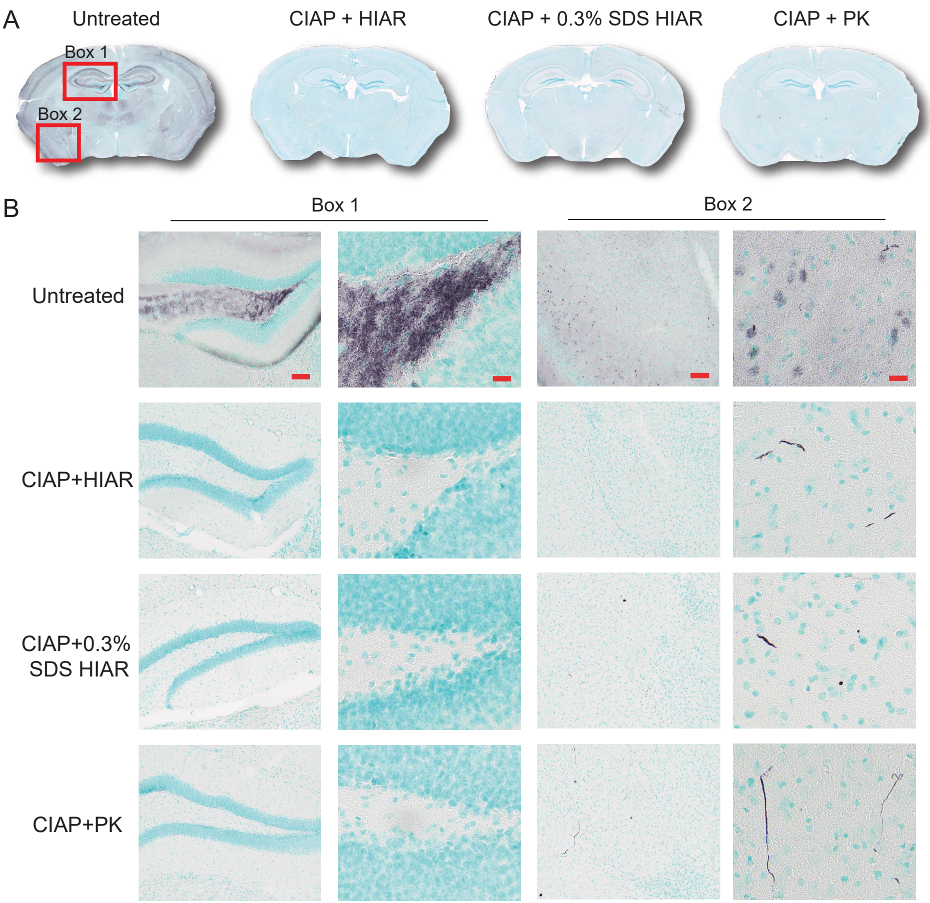


Supplementary Figure 2

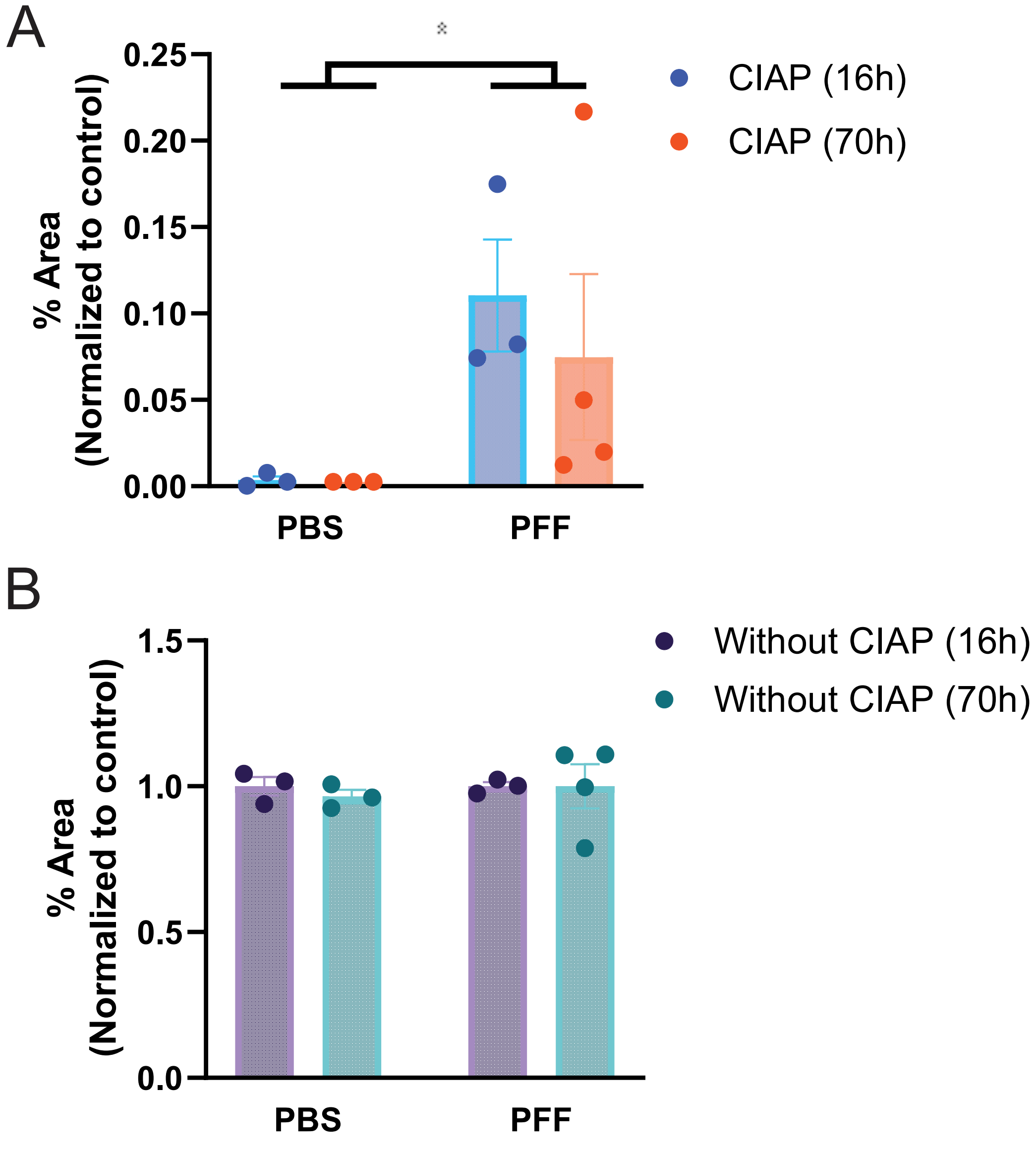


Supplementary Figure 3

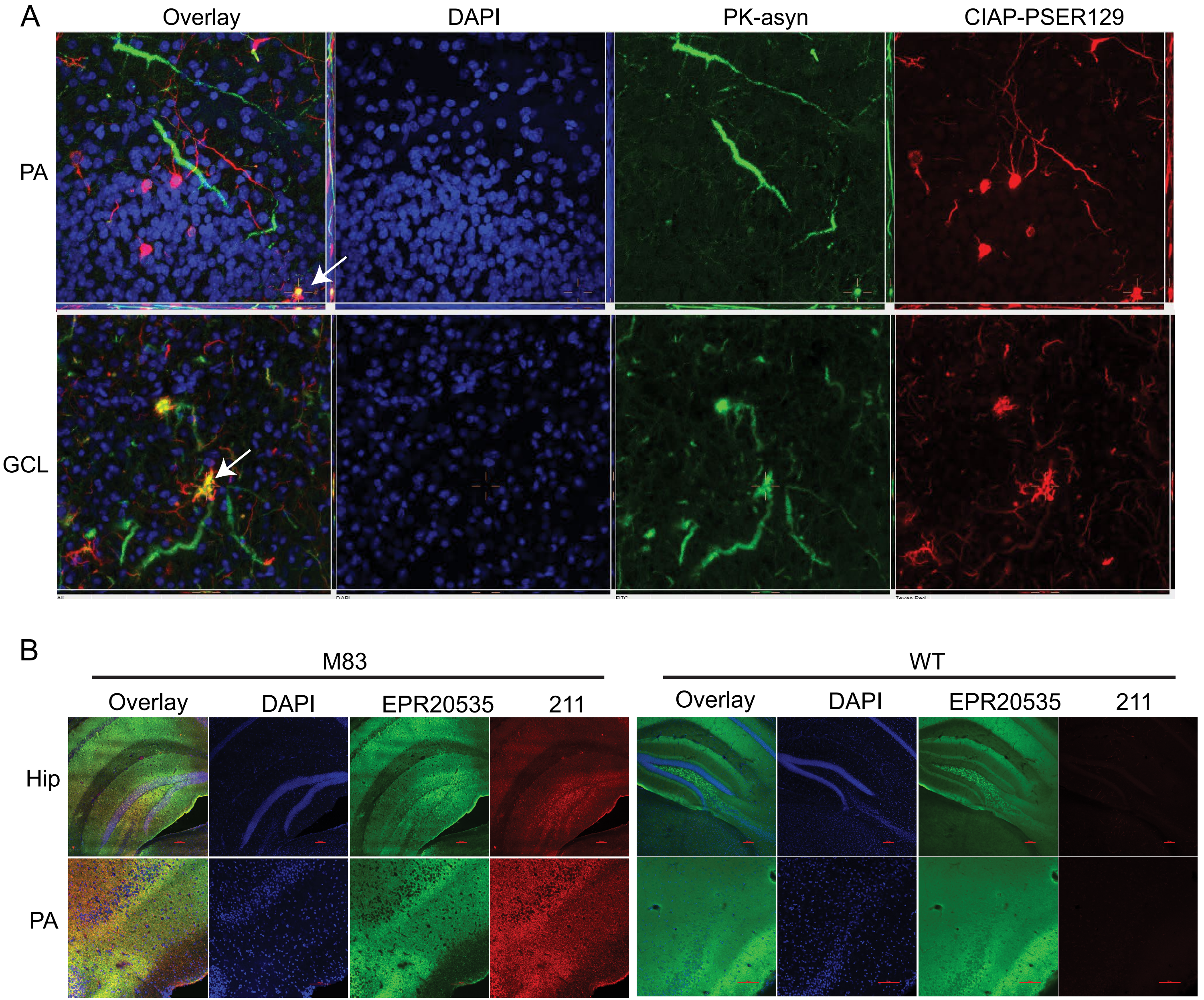


Supplementary Figure 4
